# Enzyme-independent functions of HDAC3 in the adult heart

**DOI:** 10.1101/2024.12.29.630635

**Authors:** Sichong Qian, Chen Zhang, Wenbo Li, Shiyang Song, Guanqiao Lin, Zixiu Cheng, Wenjun Zhou, Huiqi Yin, Haiyang Li, Hu-Ying Shen, Zheng Sun

**Affiliations:** Department of Cardiovascular Surgery, Beijing Anzhen Hospital, Capital Medical University, Beijing, China; Department of Medicine – Endocrinology, Baylor College of Medicine, Houston, Texas, USA; Division of Cardiothoracic Surgery, Michael E. DeBakey Department of Surgery, Baylor College of Medicine, Houston, Texas, USA; Children’s Heart Center, Institute of Cardiovascular Development and Translational Medicine, The Second Affiliated Hospital and Yuying Children’s Hospital of Wenzhou Medical University, China; Department of Molecular and Cellular Biology, Baylor College of Medicine, Houston, Texas, USA

## Abstract

The cardioprotective effects of histone deacetylase (HDAC) inhibitors (HDIs) are at odds with the deleterious effects of HDAC depletion. Here, we use HDAC3 as a prototype HDAC to address this contradiction. We show that adult-onset cardiac-specific depletion of HDAC3 in mice causes cardiac hypertrophy and contractile dysfunction on a high-fat diet (HFD), excluding developmental disruption as a major reason for the contradiction. Genetically abolishing HDAC3 enzymatic activity without affecting its protein level does not cause cardiac dysfunction on HFD. HDAC3 depletion causes robust downregulation of lipid oxidation/bioenergetic genes and upregulation of antioxidant/anti-apoptotic genes. In contrast, HDAC3 enzyme activity abolishment causes much milder changes in far fewer genes. The abnormal gene expression is cardiomyocyte-autonomous and can be rescued by an enzyme-dead HDAC3 mutant but not by an HDAC3 mutant (Δ33-70) that lacks interaction with the nuclear-envelope protein lamina-associated polypeptide 2β (LAP2β). Tethering LAP2β to the HDAC3 Δ33-70 mutant restored its ability to rescue gene expression. Finally, HDAC3 depletion, not loss of HDAC3 enzymatic activity, exacerbates cardiac contractile functions upon aortic constriction. These results suggest that the cardiac function of HDAC3 in adults is not attributable to its enzyme activity, which has implications for understanding the cardioprotective effects of HDIs.

## INTRODUCTION

Histone deacetylase (HDAC) inhibitors (HDIs) are cardioprotective in a growing list of preclinical animal models. HDIs that inhibit all classical zinc-dependent HDACs are known as pan-HDIs. Pan-HDI Trichostatin A (TSA) attenuates cardiac hypertrophy and preserves contractile functions in transverse aortic constriction (TAC) or angiotensin II-induced hypertrophy animal models^1–3^. TSA is also protective against myocardial infarction-induced contractile dysfunction ^4,5^. Suberoylanilide hydroxamic acid (SAHA), another pan-HDI, reduces myocardial fibrosis in the TAC model^6^ and attenuates myocardial injury in the isoproterenol model^7^. Givinostat (also known as ITF2357), another pan-HDI, improves diastolic function in the uninephrectomy (UNX)/deoxycorticosterone acetate (DOCA)-induced diastolic dysfunction model and the heart failure with preserved ejection fraction (HFpEF) model in Dahl salt-sensitive rats^8,9^. Mocetinostat, a benzamide-based inhibitor of Class I HDACs, improves contractile functions after myocardial infarction^10^ and attenuates cardiac hypertrophy in the TAC model^11^. However, the mechanism underlying HDIs-mediated cardiac benefits is not completely understood.

Opposite to the beneficial effects of HDIs in the heart, genetic mouse models with the whole-body or cardiac-specific knockout of HDACs often show cardiac defects ^12^. The 11 classical zinc-dependent HDACs in mammals can be grouped into classes depending on their sequence homology: Class I HDACs (HDAC1, HDAC2, HDAC3, and HDAC8); Class IIa HDACs (HDAC4, HDAC5, HDAC7, and HDAC9); Class IIb HDAC (HDAC6 and HDAC10); and Class V HDAC (HDAC11). Knockout of both HDAC1 and HDAC2 in the heart results in neonatal lethality, accompanied by cardiac arrhythmias and dilated cardiomyopathy, while deletion of HDAC1 or HDAC2 individually does not lead to obvious phenotype ^13^. Knockout of HDAC3 caused cardiac hypertrophy and affected lipid metabolism ^14,15^. HDAC3 is also essential for epicardial development ^16^. The mice with inactivated HDAC5 or HDAC9 were sensitive to stress signals, such as pressure overload and calcineurin stimulation ^17,18^.

Thus, the beneficial effects of HDIs in preclinical heart disease models are at odds with the deleterious effects of HDAC depletion in genetic mouse models. We chose to focus on HDAC3 to address this question for several reasons. (1) HDAC3 is responsible for the enzyme activity of Class IIa HDACs. Class IIa HDACs have low intrinsic HDAC enzymatic activities due to a histidine substitution on the key catalytic tyrosine site^19^. As a result, most catalytic activity of Class IIa HDACs is attributed to HDAC3, which exists in the same multiprotein corepressor complexes ^20^. (2) HDAC3 has strong enzyme activity and is considered a primary target of many HDIs. (3) Knockout of HDAC3 from different developmental stages generates different cardiac phenotypes. Knockout of HDAC3 in the heart at around embryonic day 9.5 using the α-myosin heavy chain (α-MHC)-Cre driver caused cardiac hypertrophy and lethal heart failure by the age of 3-4 months on a normal chow diet^14^. In contrast, knockout of HDAC3 in the heart postnatally using the muscle creatine kinase (MCK)-Cre driver does not have drastic cardiac phenotype on normal chow diet even at the age of over a year, and only caused cardiac hypertrophy and lethal heart failure when mice were fed with high-fat diet (HFD)^15^. These results suggest the developmental effects could account for the HDIs vs. HDAC deletion differences.

The enzyme activity of HDAC3 relies on forming a stable protein complex with nuclear receptor corepressor (NCoR or NCOR1) or silencing mediator for retinoid and thyroid receptor (SMRT or NCOR2). The deacetylation activation domain (DAD) in NCoR/SMRT binds to HDAC3 and causes a conformational change of HDAC3 protein, which makes the catalytic channel accessible to the substrate ^21,22^. Purified HDAC3 protein does not show much enzyme activity by itself but gains robust enzyme activity after adding in purified DAD ^22,23^. Conversely, whole-body knock-in of missense mutations in the DAD of NCoR/SMRT in mice (NS-DADm mice) abolishes the enzyme activity of HDAC3 without affecting its protein levels in multiple tissues ^24^. Here, we address the function of HDAC3 in the adult heart and its functional reliance on enzyme activity.

## RESULTS

### Inducible HDAC3 depletion in adult hearts causes contractile dysfunctions with high-fat-feeding

One obvious explanation for the contradiction between the beneficial effects of HDIs and the deleterious effects of HDAC depletion is the differential effects on development. Conditional knockout of an HDAC can disrupt cardiac development, while the beneficial effects of HDIs were mainly observed in adult mice. Therefore, we first address whether the previously reported detrimental cardiac effects from HDAC3 knockout are due to disruption of the cardiac developmental processes. We previously reported postnatal depletion of HDAC3 in the heart using MCK-Cre ^15^. However, the cardiac phenotype from this mouse model can still be confounded by the potential disruption of postnatal cardiac development. To eliminate the developmental effects, we crossbred HDAC3 floxed mice (HDAC3^loxP/loxP^)^15^ with cardiac-specific tamoxifen-inducible MerCreMer driver (αMHC-MerCreMer, JAX #005657)^25^ to generate HDAC3^loxP/loxP^/αMHC-MerCreMer mice for inducible knockout (referred to as “iKO**”**). Tamoxifen was administered at 7 weeks old in both iKO and the αMHC-MerCreMer mice as the wild-type control (WT). We confirmed the efficient depletion of HDAC3 by western blot analyses (**Fig 1A**). The iKO mice showed differential expression of metabolic genes on normal chow (**Fig 1B**), similar to the HDAC3^f/f^/MCK-Cre mice, as we previously reported^15^. The iKO mice did not show cardiac defects on normal chow, which is expected from normal cardiac functions in the HDAC3^loxP^/MCK-Cre line (referred to as “KO”)^15^. Therefore, we fed iKO mice with a high-fat diet (HFD) starting at 7 weeks old. HFD for 2-3 months does not induce robust cardiac hypertrophy or systolic dysfunction in WT mice, as expected from previous studies^26,27^. The iKO mice showed normal body weight gain on HFD (**Fig 1C**) but enlarged hearts at 16 weeks old (**Fig 1D**). Myocardial gene expression of atrial natriuretic peptide (ANP) and B-type natriuretic peptide (BNP) was significantly elevated in iKO mice (**Fig 1E**). iKO heart showed bigger cardiomyocytes (**Fig 1F-G**) and widespread fibrosis in the heart (**Fig 1H-I**). Echocardiography demonstrated that iKO mice developed severe cardiac hypertrophy at 4 months old after 2 months on HFD, with significant left ventricular wall thickening compared to WT mice (**Fig 1J-K**). The iKO mice developed systolic dysfunction, as evidenced by impaired ejection fraction (EF) and fractional shortening (FS) (**Fig 1K**). Thus, inducible depletion of cardiac HDAC3 in adult mice on top of 8-9 weeks of HFD feeding caused comparable cardiac dysfunction as postnatal HDAC3 depletion in combination with a similar duration of HFD feeding ^15^. Therefore, the deleterious effects of HDAC3 depletion in the heart are not due to development disruption.

**Figure 1.**
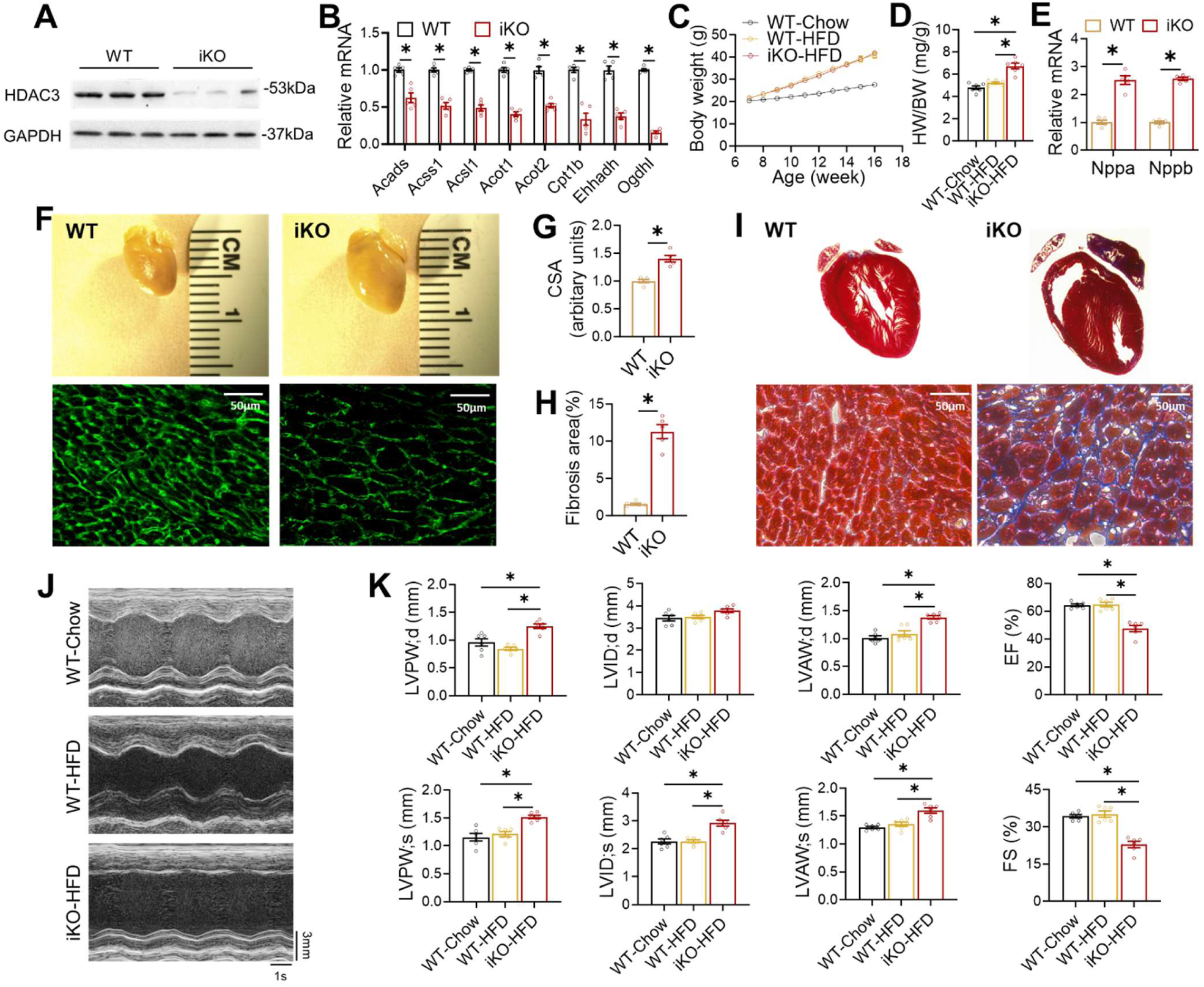
Inducible HDAC3 depletion in adult hearts impairs contractile functions on a high-fat diet (HFD). (**A**) Western blot analysis of hearts from 4-month-old mice after tamoxifen injection. (**B**) RT-qPCR analysis of the heart from 4-month-old mice on chow diet. n = 5. (**C**) Body weight (BW) on HFD or chow diet. HFD started at 7 weeks old. n = 6. (**D**) Heart weight (HW) to body weight (BW) ratio of 4-month-old mice on HFD or chow diet. n = 6. (**E**) RT-qPCR analysis of myocardial ANP and BNP from 4-month-old mice on HFD. (**F**) Gross pictures and wheat germ agglutinin (WGA) staining of hearts from 4-month-old mice on HFD. Scale bar: 50 µm. (**G**) WGA quantification of cardiomyocyte cross-sectional area. n = 5. (**H**) Percentage of fibrosis area in trichrome staining transversal sections. n = 5. (**I**) Trichrome staining of hearts from 4-month-old mice on HFD. Scale bar: 50 µm. (**J**) Representative M-mode recordings of mouse hearts in echocardiography. (**K**) Echocardiography of geometry and systolic functions in 15-weeks-old mice on HFD, n = 6. All data are mean ± S.E.M. * p < 0.05 by t-test or one-way ANOVA with Holm-Sidak’s post hoc.

### Abolishing HDAC3 enzymatic activity does not cause cardiac defects on HFD

We next sought to address the role of HDAC3 enzyme activity in the heart without disrupting the HDAC3 protein level. The deacetylation activation domain (DAD) in NCoR/SMRT binds to HDAC3 and causes a conformational change of HDAC3 protein, making the catalytic channel accessible to the substrate ^21,22^. Purified HDAC3 protein does not show much enzyme activity by itself but gains robust enzyme activity after adding in purified DAD ^22,23^. Conversely, the whole-body knock-in NS-DADm mouse line harbors homozygous mutations in the DAD of both NCoR and SMRT (NCoR-Y478A and SMRT-Y470A) showed normal HDAC3 protein levels, but ablated HDAC3 enzymatic activity in multiple tissues ^24^. Therefore, we used the NS-DADm mice to address the role of HDAC3 enzyme activity in the heart. Since iKO mice have a similar phenotype as the HDAC3^loxP^/MCK-Cre (KO) mice, we used KO mice to avoid potential complications from tamoxifen injection.

To measure the deacetylase enzyme activity of HDAC3 on any target, not just histone targets, we used HDAC3-specific antibodies to immunoprecipitate HDAC3 from the total protein lysates of the heart and subjected the immunoprecipitates to western blot analysis and a peptide-based enzyme activity assay (**Fig 2A-B**). While HDAC3 protein levels remain normal in the adult NS-DADm heart, the interaction between HDAC3 and NCOR1 or TBL1XR1, another stable component of the NCOR complex, was disrupted (**Fig 2A**). In addition, the HDAC3 deacetylase activity was abolished in the HDAC3 immunoprecipitates from the NS-DADm heart, (**Fig 2B**). NS-DADm mice gained similar body weight on HFD as WT and KO mice (**Fig 2C**). As we have reported before ^15^, the KO heart was markedly enlarged compared to WT after feeding HFD for 9 weeks (**Fig 2D**-**E**), with enlarged cardiomyocyte size (**Fig 2E-F**), widespread interstitial fibrosis (**Fig 2E** and **2G**), and elevated myocardial expression of ANP and BNP (**Fig 2H**). Echocardiography showed that KO mice developed cardiac hypertrophy and systolic dysfunction (**Fig 2I-J**). In contrast, NS-DADm mice showed no defects after HFD (**Fig 2D**-**J**). These data demonstrate that abolishing HDAC3 enzymatic activity without affecting its protein levels does not affect cardiac functions on HFD. Thus, the detrimental effects of HDAC3 depletion on contractile functions in the presence of HFD are not due to the abolishment of HDAC3 enzyme activity.

**Figure 2.**
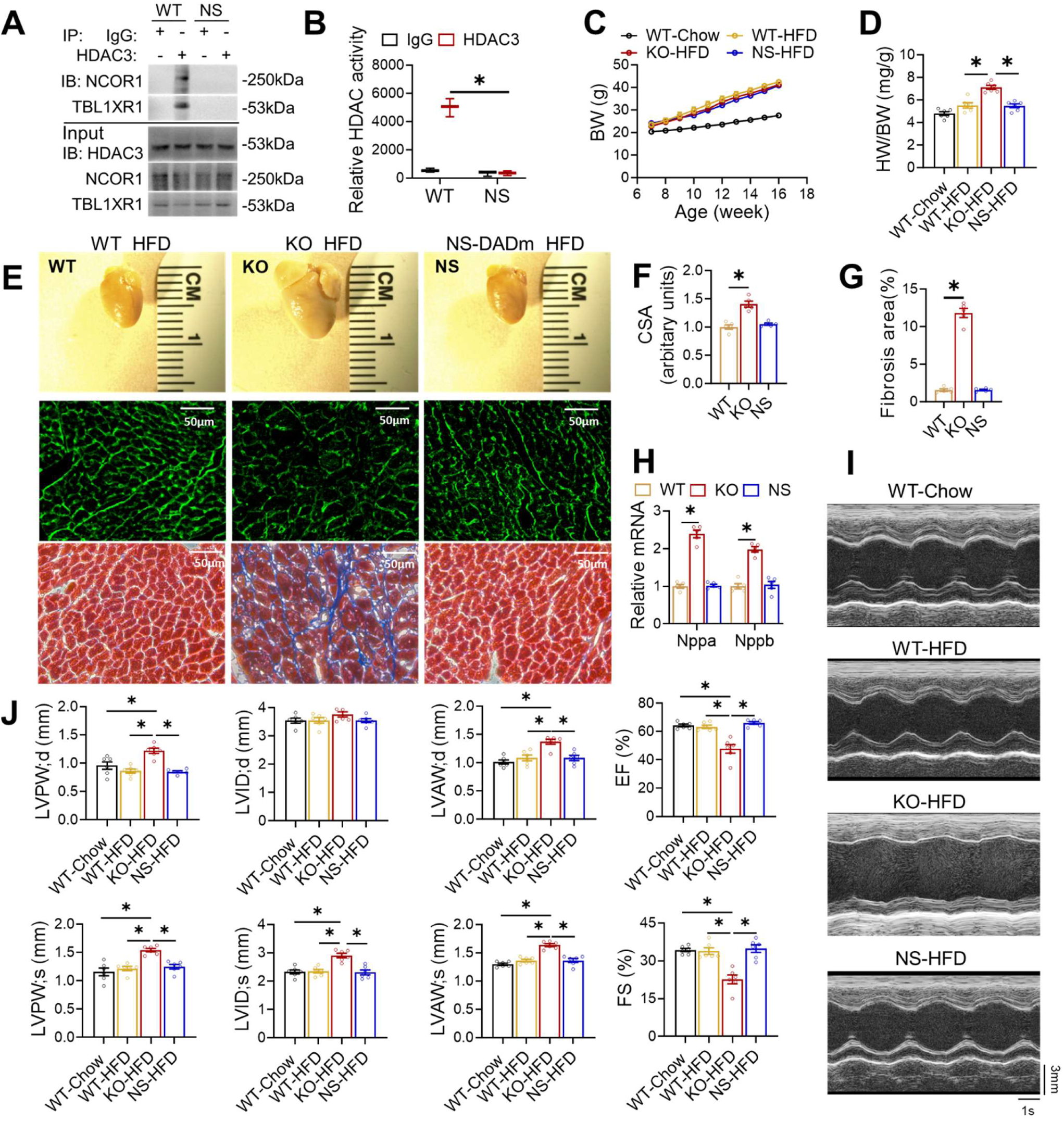
Abolishing HDAC3 enzymatic activity does not cause cardiac defects on HFD. (**A**) Immunoblot (IB) analysis of immunoprecipitates (IP) and input lysates from hearts of 4-month-old WT and NS-DADm mice with the indicated antibodies. (**B**) Fluorescence-based HDAC enzyme assay using lysates of mice hearts after immunoprecipitation with HDAC3 antibody or normal IgG. n =3. (**C**) Body weight (BW) on HFD or chow diet. HFD started at 7 weeks old. n = 6. (**D**) Heart weight (HW) to body weight (BW) ratio of 4-month-old mice on HFD or chow diet. n = 6. (**E**) Gross pictures, wheat germ agglutinin (WGA) staining, and trichrome staining of hearts from 4-month-old mice fed on HFD. Scale bar: 50 µm. (**F**) WGA quantification of cardiomyocyte cross-sectional area on samples from HFD-fed mice. n = 5. (**G**) Percentage of fibrosis area in trichrome staining transversal sections. n = 5. (**H**) RT-qPCR analysis of the myocardial ANP and BNP from 4-month-old mice fed on HFD. n = 5. (**I**) Representative M-mode recordings of mouse hearts in echocardiography of 4-month-old mice. (**J**) Echocardiography of geometry and systolic functions of 4-month-old mice, n = 6. All data are mean ± S.E.M. * *p* < 0.05 by t-test or one-way ANOVA with Holm-Sidak’s post hoc.

### Distinct transcriptomic changes between HDAC3 depletion and loss of HDAC3 enzyme activity

To address whether differential cardiac effects between HDAC3 depletion and loss of HDAC3 enzyme activity is due to differential transcriptomic effects, we performed RNA-seq analyses in the hearts of KO mice and NS-DADm mice. We harvested hearts at 6 weeks old on normal chow to exclude potential confounding effects of contractile dysfunction on cardiac gene expression. There were about 17 times more differentially expressed genes (DEGs) in the KO vs. WT hearts than in the NS-DADm vs. WT hearts (**Fig 3A**). The downregulated DEGs (KO vs. WT) were enriched in mitochondrial fatty acid metabolism (**Fig 3B**), while the upregulated DEGs (KO vs. WT) were enriched in cell survival and antioxidant response (**Fig 3C**). Most downregulated metabolic genes were not significantly altered in the NS-DADm vs. WT control (**Fig 3D**-**E**), which were further confirmed by RT-qPCR analyses (**Fig 3F**). Both KO and NS-DADm hearts showed upregulation of genes involved in cell survival and antioxidant response compared to their respective WT controls, but the fold-changes in the NS-DADm vs. WT comparison were much less than those in the KO vs. WT comparison (**Fig 3G**-**I**). The lack of differential expression in metabolic genes in NS-DADm is in keeping with the lack of contractile dysfunctions in NS-DADm. These results suggest that HDAC3 uses an enzyme-independent mechanism to regulate lipid metabolism, which could contribute to the detrimental effects of HDAC3 depletion. We do not suggest that these lipid metabolic genes are direct target genes of HDAC3, as this question is not relevant to whether HDAC3 function requires its deacetylase enzyme activity.

**Figure 3.**
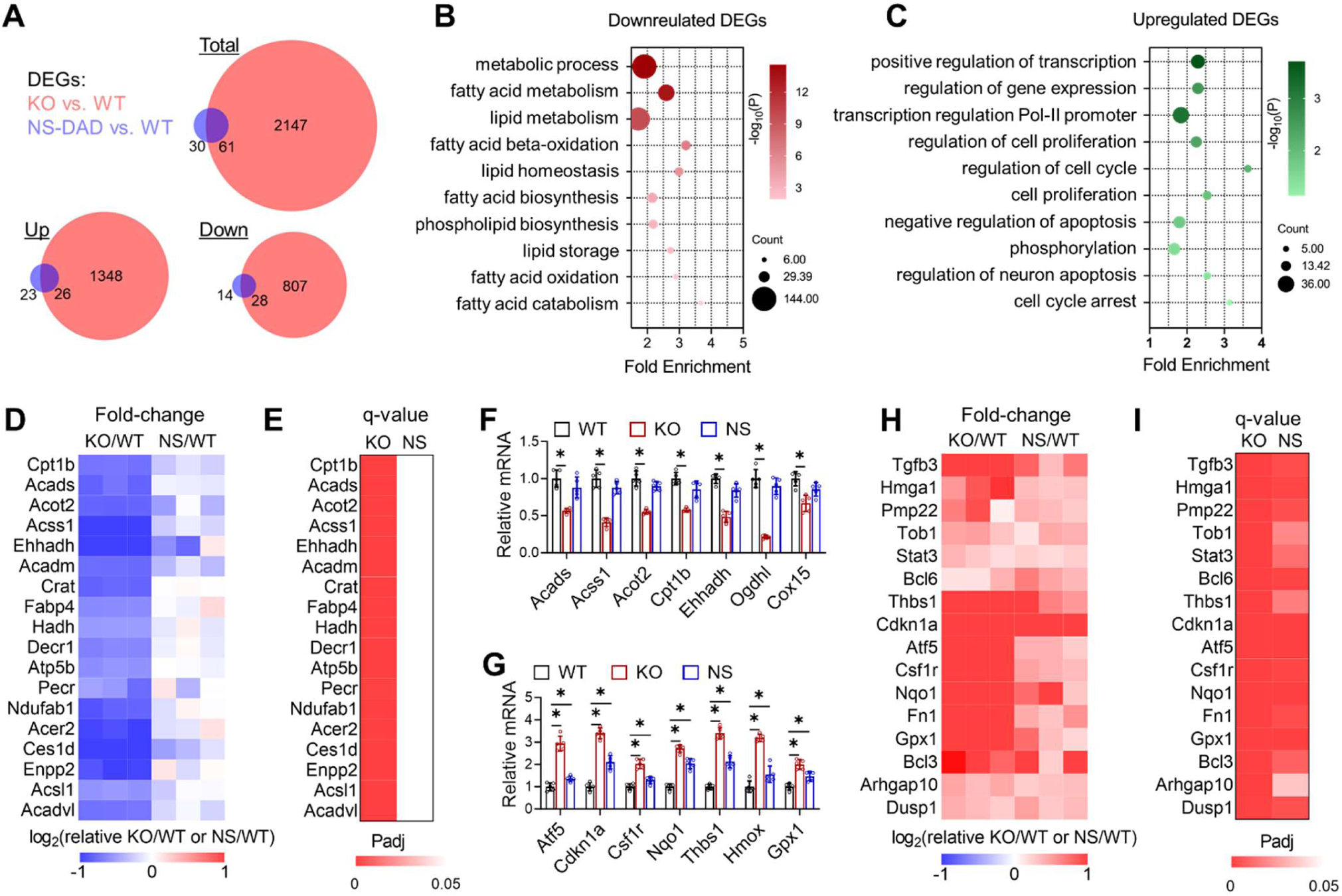
Enzyme-independent regulation of fatty acid oxidation genes by HDAC3 in cardiomyocytes. (**A**) Overlap of differentially expressed genes (DEGs) in the NS-DADm (NS) and KO hearts compared to their respective WT controls on normal chow of 6-week-old mice, identified by RNA-seq. DEGs cutoff: q<0.05 and |log2Fold-Change| >1. (**B**) Gene ontology analysis of the pooled downregulated genes (KO vs. WT and NS-DADm vs. WT, q<0.05). (**C**) Gene ontology analysis of the pooled upregulated genes (KO vs. WT and NS-DADm vs. WT, q<0.05). (**D**) A heatmap of fold-change top-downregulated genes in the KO/WT and NS-DADm/WT comparisons. (**E**) A heatmap of adjusted P values (Padj) in the KO/WT and NS-DADm/WT comparisons. (**F-G**) RT-qPCR analysis of mRNA from the heart of 6 weeks mice. n = 5. * *p* < 0.05 by one-way ANOVA with Holm-Sidak’s post hoc. (**H**) A heatmap of fold-change top-upregulated genes in the KO/WT and NS-DADm/WT comparisons. (**I**) A heatmap of Padj values in the KO/WT and NS-DADm/WT comparisons.

### Discrete cell-autonomous effects between HDAC3 depletion and HDAC enzyme inhibition

Considering that many systemic or paracrine factors could contribute to the gene expression changes *in vivo*, we sought to address whether the differential effects of HDAC3 depletion and HDAC3 enzyme inactivation on gene expression and metabolism are cell-autonomous in an *in vitro* cell culture model. We first examined the effects of HDAC3 deletion in AC16 cardiomyocytes. Western blot analysis confirmed the efficient depletion of HDAC3 by adenovirus-mediated delivery of single-guiding RNAs (sgRNA) and Cas9 targeting HDAC3 (**Fig 4A**). Global histone acetylation levels were not robustly altered by knocking out a single HDAC, which is consistent with previous studies^28,29^. RT-qPCR analysis demonstrated that HDAC3 depletion in AC16 cells downregulated genes in fatty acid oxidation and upregulated genes in anti-apoptosis and antioxidant responses (**Fig 4B**-**C**). The metabolic flux analysis with the ^3^H-palmitate isotope tracer showed that HDAC3 depletion reduced the fatty acid oxidation rate (**Fig 4D**). These results suggest that the effects of HDAC3 depletion on gene expression and lipid metabolism can be recapitulated in cultured cardiomyocytes.

**Figure 4.**
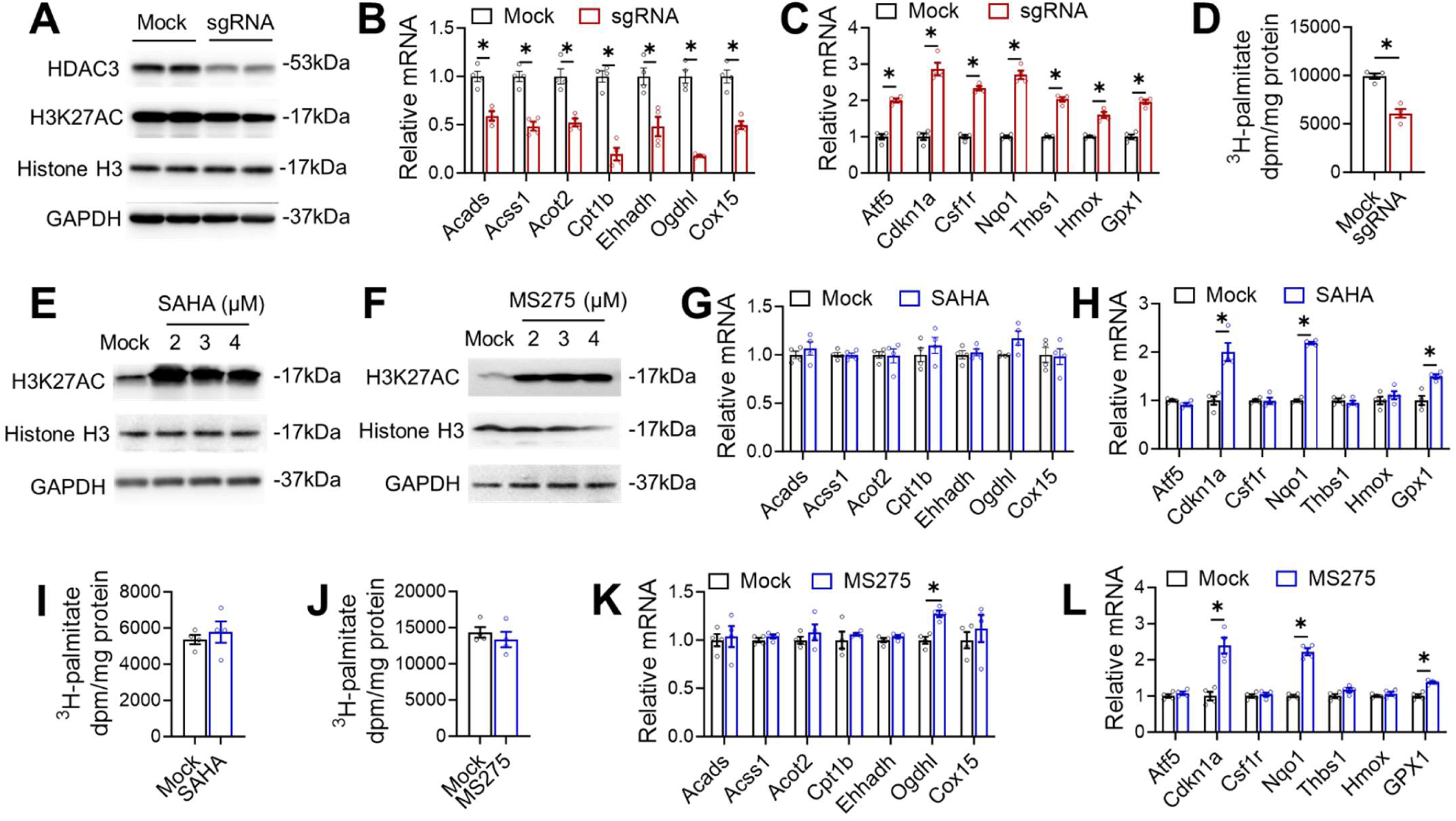
Discrete cell-autonomous effects between HDAC3 depletion and HDAC enzyme inhibition. (**A**) Western blot analysis of lysates from AC16 cells with HDAC3 knock-down. (**B-C**) RT-qPCR analysis of the AC16 cells with HDAC3 knock-down. n = 4. (**D**) Fatty acid oxidation (FAO) of the AC16 cells with HDAC3 knock-down. n = 4. (**E-F**) Western blot analysis of lysates from AC16 cells administrated with 2 uM MS275 or SAHA at the indicated dosages, along with vehicle control (mock). (**G-H**) RT-qPCR analysis of the AC16 cells treated with 2 uM SAHA or MS275. (**I-J**) Fatty acid oxidation (FAO) of the AC16 cells treated with 2 uM SAHA or MS275. (**K-L**) RT-qPCR analysis of the AC16 cells treated with 2 uM SAHA or MS275. All data are mean ± S.E.M. * *p* < 0.05 by t-test.

We next examined the effects of HDIs, such as suberoylanilide hydroxamic acid (SAHA) and entinostat (MS-275), in AC16 cells. Despite increased global histone acetylation (**Fig 4E**-**F**), SAHA and MS-275 did not affect the expression of lipid oxidation genes (**Fig 4G-H**) and did not alter the fatty acid oxidation rate (**Fig 4I**-**J**). These results are consistent with the lack of significant changes in fatty acid oxidation genes *in vivo* in NS-DADm mice. HDIs also upregulated some of the genes involved in cell survival and antioxidant responses in AC16 cells (**Fig 4K**-**L**), which is consistent with the upregulation of these genes in the NS-DADm vs. WT hearts. We performed similar western blot and RT-qPCR analyses in induced pluripotent stem cell (iPSC)-derived cardiomyocytes (iPSC-CM) and observed similar results (**Supplemental Fig S1**). These results demonstrated that the differential effects of HDAC3 depletion vs. HDAC3 enzyme inhibition are cardiomyocyte-autonomous.

### HDAC3-LAP2β interaction in the enzyme-independent function of HDAC3 in cardiomyocytes

The enzyme-dependent repression of cell survival and antioxidant genes by HDAC3 is generally in line with the canonical view of how HDACs repress gene transcription through histone deacetylation and chromatin remodeling ^30,31^, although some of these genes could be indirectly altered due to the altered expression of the direct HDAC3 target genes. Rather than dissecting the direct vs. indirect targets of HDAC3 enzyme-dependent target genes, we want to focus on the enzyme-independent target genes because the mechanism is more intriguing. Recent studies suggest that the interaction of HDAC3 with an inner nuclear membrane protein LAP2β is required for the enzyme-independent function of HDAC3 in restricting the precocious cardiac progenitor differentiation ^32^, and LAP2 dysfunction protein is associated with abnormal lipid metabolism in the hepatocyte ^33^. Therefore, we wonder whether a similar mechanism is responsible for the enzyme-independent function of HDAC3 in lipid metabolism in mature cardiomyocytes.

Previous studies indicated the interaction between HDAC3 and LAP2β was mediated by the 38-amino acid domain of HDAC3 (33-70) ^32^. We confirmed that Flag-tagged HDAC3 wild-type (WT) interacts with HA-tagged LAP2β, and a deletion mutant of HDAC3 (Δ33-70) abolished such interaction (**Fig 5A**) without affecting interaction with endogenous NCOR1 or TBL1XR1 (**Fig 5B**). In comparison, the missense mutation of HDAC3 on the catalytic site (Y298H or YH) retained interaction with LAP2β (**Fig 5A**) but abolished the enzyme activity (**Fig 5C**). Another missense mutation of HDAC3 (K25A or KA) from our previous report ^29^ disrupted interaction with NCOR1/TBL1XR1 and served as a negative control for the co-immunoprecipitation assay (**Fig 5B**). LAP2β still binds to the HDAC3 K25A mutant (**Supplemental Fig S2A**), suggesting that the HDAC3-LAP2β interaction is independent of the HDAC3-NCOR interaction and thus likely remains intact in NS-DADm mice. HDAC inhibitors. SAHA or MS-275 did not affect the binding of HDAC3 with LAP2β (**Supplemental Fig S2B**). We also fused HDAC3 WT or Δ33-70 to LAP2β (WT-L and Δ33-70-L) (**Fig 5A-C**) and used these mutants in a rescue experiment on top of HDAC3 depletion in AC16 cells. All HDAC3 constructs were engineered to evade the sgRNA targeting regions. We used adenovirus vectors to deliver HDAC3 constructs into AC16 cells, which can efficiently infect nearly all cells in the culture (**Fig 5D**). The virus dosage was adjusted so that all exogenous HDAC3 was expressed at a similar level as the endogenous HDAC3 (**Fig 5E**).

**Figure 5.**
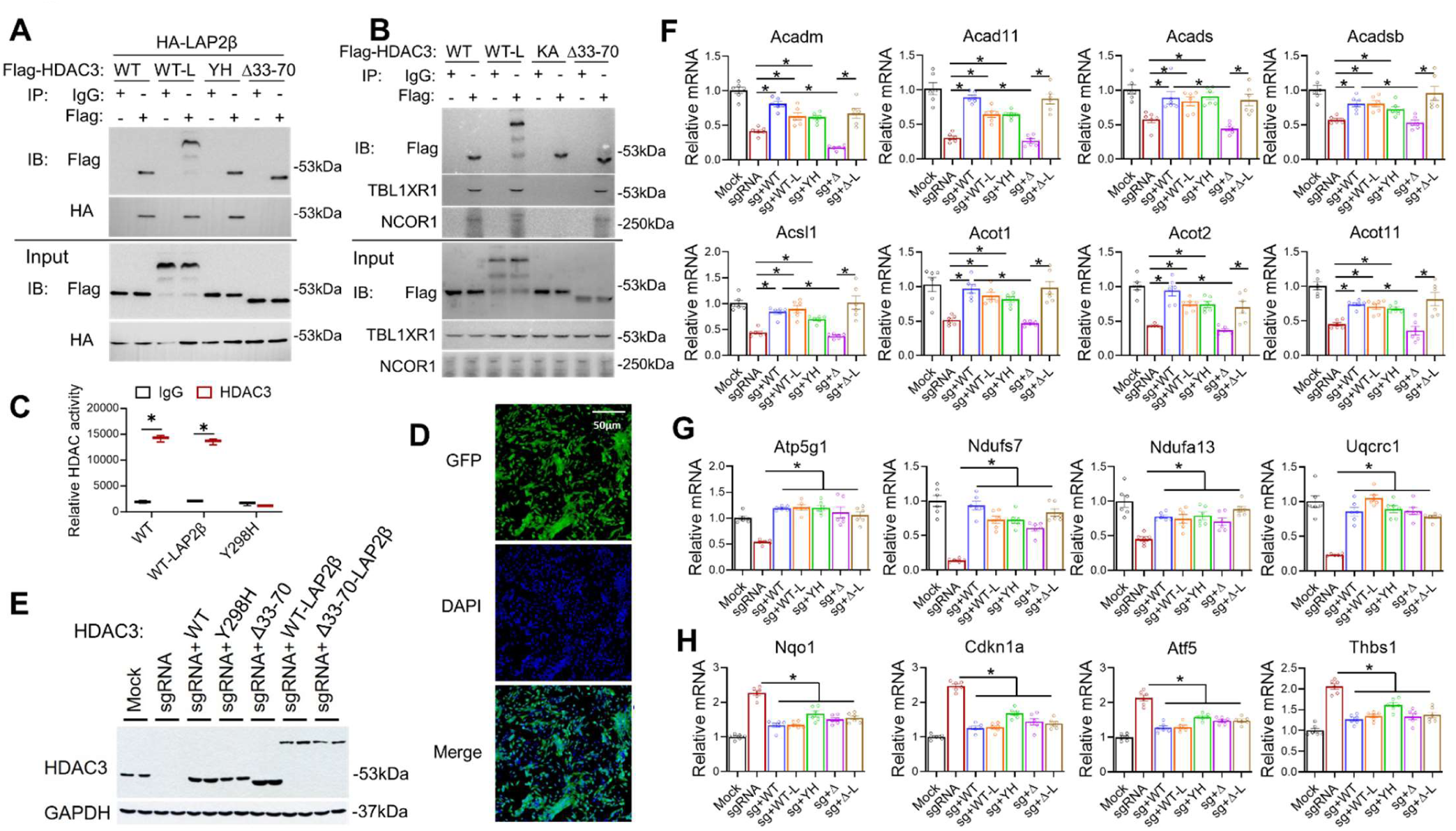
HDAC3-LAP2β interaction in the enzyme-independent function of HDAC3 in cardiomyocytes. (**A-B**) Immunoblot (IB) analysis of immunoprecipitates (IP) and input lysates from HEK293 cells transfected with indicated plasmids with the indicated antibodies. (**C**) Fluorescence-based HDAC enzyme assay using lysates of HEK293 cells transfected with indicated constructs after immunoprecipitation with HDAC3 antibody or normal IgG, n = 3. (**D**) Fluorescence microscopy of AC16 cells infected with adenovirus expressing GFP. (**E**) Western blot analysis of AC16 cells infected with the indicated adenovirus vectors. (**F-H**) RT-qPCR analysis of the AC16 cells infected with the indicated adenovirus vectors. n = 4. All data are mean ± S.E.M. * *p* < 0.05 by t-test or one-way ANOVA with Holm-Sidak’s post hoc.

Re-expression of HDAC3 WT in AC16 cells rescued KO-induced changes in lipid oxidation gene expression (**Fig 5F**). The enzyme-dead Y298H mutant behaved similarly to WT in almost all genes tested (**Fig 5F-H**). These results are consistent with the *in vivo* results in NS-DADm mice and the *in vitro* data with HDIs. These results demonstrated that HDAC3-mediated regulation of gene expression in cardiomyocytes is independent of HDAC3 enzyme activity. The HDAC3 Δ33-70 mutant did not rescue lipid oxidation gene expression (**Fig 5F**), but artificially tethering LAP2β to the HDAC3 Δ33-70 mutant regained the ability to rescue lipid oxidation gene expression. Consistent with the gene expression results, the fatty acid oxidation assay showed a similar pattern of rescue (**Supplemental Fig S2C**). These results suggest that the interaction with LAP2β is crucial for the HDAC3-mediated regulation of lipid oxidation. Notably, HDAC3 Δ33-70 mutant rescued some oxidative phosphorylation genes (**Fig 5G**) and genes involved in cell survival and antioxidant responses (**Fig 5H**) as well as HDAC3 WT, suggesting that the regulation of these genes by HDAC3 does not require LAP2β. In summary, HDAC3 modulates gene expression in cardiomyocytes through an enzyme-independent mechanism. Its interaction with LAP2β contributes significantly to the regulation of downstream target genes, although not all of them are influenced by this interaction. It is unclear what caused such gene-specific regulation and whether the interaction with LAP2β is the major contributor to the in vivo function of HDAC3.

### Distinction between HDAC3 depletion and activity suppression upon pressure overload

We have used HFD-induced obesity as a pathological stressor to study HDAC3 enzyme-dependent and - independent functions. To address whether the insights obtained from the HFD model are generalizable, we sought to use transverse aortic constriction (TAC), a common model for pressure overload-induced cardiac hypertrophy and heart failure. We subjected WT, KO, and NS-DADm mice to TAC at 7-8 weeks old. All groups of mice had similar body weights (**Fig 6A**). After TAC, KO mice, but not NS-DADm mice, showed more prominent cardiac hypertrophy than WT mice at 12 weeks old (**Fig 6B-G**). Specifically, KO mice after TAC showed enlarged hearts (**Fig 6B** and **6G**), enlarged cardiomyocytes (**Fig 6B-C**), prominent interstitial fibrosis (**Fig 6D-E**), and elevated ANP and BNP gene expression in the heart compared to WT (**Fig 6F**). Echocardiography analysis showed that KO mice, but not NS-DADm mice, displayed cardiac dilation and severe systolic dysfunction, as evidenced by severe ventricular wall and reduced ejection fraction (**Fig 6H-I**). These findings suggest that the adverse effects of HDAC3 depletion on cardiac contractile functions are independent of its enzyme activity. Abolishing HDAC3 enzyme activity per se, without altering its protein levels, does not increase susceptibility to overload-induced hypertrophy or contractile dysfunctions.

**Figure 6.**
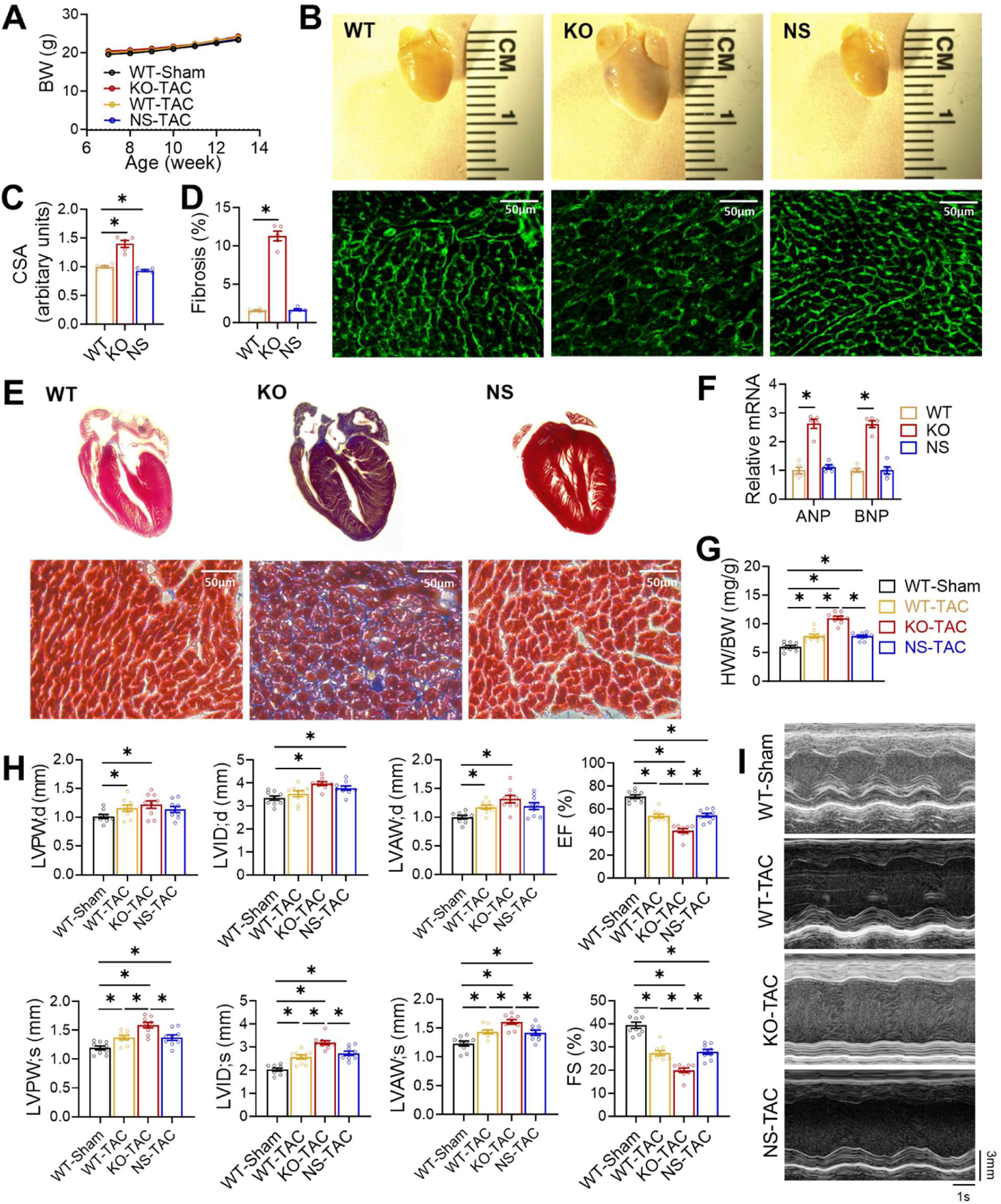
Distinct cardiac outcomes between HDAC3 depletion and activity suppression upon pressure overload. (**A**) Body weight (BW) after transverse aortic constriction (TAC) or sham surgery at 2-month-old. n = 9. (**B**) Gross pictures and wheat germ agglutinin (WGA) staining of hearts from 3-month-old mice after TAC. Scale bar: 50 µm. (**C**) WGA quantification of cardiomyocyte cross-sectional area. n = 5. (**D**) Percentage of fibrosis area in trichrome staining transversal sections. n = 5. (**E**) Trichrome staining of cross-sections of hearts from 3-month-old mice after TAC. Scale bar: 50 µm. (**F**) RT-qPCR analysis of the myocardial ANP and BNP from 3-month-old mice after TAC. n = 5. (**G**) Heart weight (HW) to body weight (BW) ratio of 3-month-old mice after TAC or sham surgery. n = 9. (**H**) Echocardiography of geometry and contractile functions of 3-month-old mice, n = 9. (**I**) Representative M-mode recordings of mouse hearts in echocardiography of 3-month-old mice. All data are mean ± S.E.M. * *p* < 0.05 by t-test or one-way ANOVA with Holm-Sidak’s post hoc.

## DISCUSSION

The dissection of enzyme-dependent and -independent functions of HDAC3 contributes to our understanding of the apparent paradox between the beneficial effects of HDIs and the deleterious effects of HDAC depletion on cardiac functions. The results suggest that HDAC3 uses an enzyme-independent, LAP2β-dependent mechanism to maintain active oxidative metabolism of the myocardium. Depletion of HDAC3 proteins represses fatty acid oxidation and compromises myocardial bioenergetics, which can contribute to contractile dysfunctions. This enzyme-independent function of HDAC3 in adult hearts is in keeping with previous studies showing the enzyme-independent function of HDAC3 in cardiac development in mice ^32,34^ and cardiac contractility in drosophila^35^. The involvement of LAP2β in lipid metabolism is in line with several recent studies suggesting the active roles of the nuclear envelope in regulating lipid metabolic genes ^33,36,37^. In particular, overexpression of a mutant LAP2 protein with disrupted interaction with nucleoplasmic lamin A caused more lipid accumulation than WT LAP2 in the hepatocytes ^33^. We also tested the alternative explanation that the adverse effect in genetic mouse models could be due to developmental disruption, which is absent in HDI treatment in adult animals. We refuted this explanation because the adult-onset HDAC3 knockout mice showed similar progressive cardiac dysfunctions as the previously reported postnatal HDAC3 knockout mice^15^.

The absence of defects in HDAC inhibitor-treated cells (compared to vehicle) or NS-DADm mice (compared to WT) suggests that the adverse cardiac outcomes of HDAC3 depletion are not attributable to the loss of its enzymatic activity. This notion is separate from whether the beneficial effects of HDAC inhibitors (compared to vehicle-treated WT) arise from HDAC3 inhibition. Since we did not see functional improvement in NS-DADm mice compared to WT, our data suggest that the beneficial effects of HDAC inhibitors are likely independent of HDAC3. Pan-HDAC inhibitors’ cardioprotective effects may be from blocking enzyme activity of other HDACs or off-target effects. For example, the HDI ITF2357 (givinostat) was shown to attenuate cardiac myofibril relaxation ^38^. SAHA was shown to regulate mitochondrial metabolism through post-translational modification of mitochondrial enzymes ^39^. The expression of mitochondrial fatty acid oxidation genes was not changed after HDI treatment ^39^, which is in line with our transcriptomic results. HDIs-mediated cardioprotection is reported to be associated with anti-apoptosis mechanisms ^30,31,40^ and mildly enhanced oxidative stress ^41^, which is consistent with the upregulated anti-apoptotic and antioxidant genes in NS-DADm hearts or after HDIs treatment in our study. However, we did not observe an improvement in cardiac functions in NS-DADm mice, probably due to complications of HDAC3 enzyme activity abolishment in other tissues.

The current study adds to a growing list of enzyme-independent functions of HDAC3. We and others have previously shown that some *in vivo* functions of HDAC3 are not dependent on its enzyme activity. For example, liver-specific knockout of HDAC3 led to hepatosteatosis and upregulated lipogenic gene expression. Mutation of the catalytic tyrosine abolished the enzyme activity of HDAC3 but can still rescue hepatosteatosis and similarly repress lipogenic gene expression as wild-type (WT) HDAC3^29^. The NS-DADm mice showed only mild hepatosteatosis and limited upregulation of lipogenic genes in the liver compared to HDAC3 depletion ^24^. These results suggest that the function of HDAC3 in liver lipid metabolism is largely independent of its enzyme activity. Similarly, the function of HDAC3 in cardiac development and spermatogenesis is not dependent on its enzyme activity, as we and others have shown ^32,42^. However, the enzyme dependency is highly tissue-specific, as the function of HDAC3 in skeletal muscle fuel metabolism is indeed through its enzymatic activity^43,44^. The function of the brain HDAC3 in neurocognition is also dependent on its enzyme activity^45,46^. Along the same line, the function of HDAC3 in B-cell development and survival requires its enzymatic activity ^47^. These results suggest that the degree to which the function of HDAC3 requires its enzyme activity is highly dependent on the tissue or cell type. Therefore, whether the HDACs function in each context requires their enzyme activity should be tested independently.

The current study has several limitations. We only characterized male mice because we did not see obvious sex differences when characterizing the original KO mice on HFD. It is intriguing that some downstream target genes are sensitive to disruptions in HDAD-LAP2β binding while others are not. We do not know the mechanism underlying the gene-specific regulation. We speculate that the chromatin context, especially what transcription factors and coregulators occupy the neighboring genomic loci, as well as the epigenomic marks and the accessibility of the loci, could all play a role in this gene-specific effect. Another possible limitation is that we did not run chromatin immunoprecipitation (ChIP) when assessing protein hyper-acetylation due to HDAC3 depletion or inhibition. ChIP is more sensitive than Western blot since it can detect loci-specific histone modifications. However, compared to the enzyme activity assay we did, the more laborious ChIP does not address non-histone targets that could be equally important for the phenotypic changes. The key scientific question of our study is not how HDAC3 regulates gene expression through histone modifications but whether HDAC3 function in the heart depends on its enzyme activity on any histone or non-histone substrate. We did not compare HDIs with genetic manipulation of HDAC3 in vivo because HDIs can have effects independent of HDAC3 or any other HDACs. In addition, HDIs administered in vivo are cleared from the body after a few hours, which is different from the continuous abolishment of HDAC3 enzyme activity in iKO or NS-DADm mice. Thus, direct comparisons between HDIs and HDAC3 iKO or NS-DADm mice for physiological outcomes could be confounded by multiple factors. Therefore, we only used HDIs in vitro to assess short-term transcriptional effects. We also did not know whether interaction with LAP2β is sufficient for the function of HDAC3 in vivo since it is technically challenging to use AAV to achieve transgene expression with a widespread and homogeneous pattern that matches the endogenous protein level. Considering that the endogenous HDAC3 protein level in the heart is relatively low, interpreting the results of such experiments could be confounded by overexpression artifacts. Finally, the current study focuses on the function of HDAC3 in cardiomyocytes in HFD or TAC pathological models. Whether HDAC3 plays a role in other cell types in these conditions is beyond the scope of the current study.

## METHODS

### Animals

HDAC3^loxP/loxP^/αMHC-MerCreMer (iKO) mice were generated from crossing HDAC3^loxP/loxP^ and transgenic αMHC-MerCreMer mice. HDAC3^loxP/loxP^ mice, HDAC3^loxP/loxP^/MCK-Cre (KO) mice, and NS-DADm mice were previously described ^15,44,46^. All mice were on the C57BL/6J genetic background. Male mice at the age of 3-4 months were used for echocardiography and histology experiments. Tamoxifen was dissolved in corn oil and injected intraperitoneally (20 mg/kg per day) for 5 consecutive days. High-fat diet (HFD) containing 60 kcal % fat was purchased from Research Diets Inc (D12492i). For euthanization, mice were placed in a chamber where carbon dioxide (CO2) was gradually introduced until the mice were unconscious and subsequently confirmed dead. All the animal care and procedures were reviewed and approved by the Institutional Animal Care and Use Committee (IACUC) at the Baylor College of Medicine and conformed to the NIH Guide for the Care and Use of Laboratory Animals.

### TAC surgery

The chronic pressure overload model was induced by transverse aorta constriction (TAC) performed on 8-week-old male mice following the established procedure^48^. After anesthetization with intraperitoneal injection of a mixture of ketamine (100 mg/kg) and xylazine (20 mg/kg), mice were intubated and placed on a respirator. Midline sternotomy was performed, and the aorta constricted at the mid-aortic arch level with a 6/0 braided silk suture using a blunted 27.5-gauge needle as a calibrator. Sham control mice underwent the same surgical procedures without constriction of the aorta.

### Echocardiography

Non-invasive transthoracic echocardiography was performed using a VisualSonics Vevo 2100 system. Two-dimensional images were obtained at 2.5-3 frames/s using a 15 MHz probe (RMV 707B, Visual Sonics) in the parasternal short-axis views to guide M-mode analysis at the midventricular level. Anesthesia was induced by 2.5% isoflurane and confirmed by a lack of response to firm pressure on one of the hind paws. During echocardiogram acquisition, isoflurane was adjusted to 1.5% to maintain a heart rate of 400-460 beats per minute. Parameters collected include: heart rate, left ventricular end-diastolic internal diameter (LVID;d), left ventricular end-systolic internal diameter (LVID;s), left ventricular end-diastolic anterior wall thickness (LVAW;d), left ventricular end-systolic anterior wall thickness (LVAW;s), left ventricular end-diastolic posterior wall thickness (LVPW;d), left ventricular end-systolic posterior wall thickness (LVPW;s), left ventricular ejection fraction (LVEF), left ventricular fractional shorting (LVFS). At the end of the procedures, all mice recovered from anesthesia without difficulties. We consistently performed left ventricular short-axis M-mode scans at the level of the bi-papillary muscles to assess LV systolic function, as well as to measure dimensions and wall thickness. To ensure the reliability of our measurements and address potential variability, we evaluated both inter-observer and intra-observer variability for all indices. This evaluation involved a blinded analysis of three randomly selected echocardiographic studies. The same observer analyzed the data on two separate occasions to assess intra-observer variability, while two independent observers analyzed the data to assess inter-observer variability. Indices of cardiac function were obtained from short-axis M-mode scans at the midventricular level, as indicated by the presence of papillary muscles in anesthetized mice. We calculated the ejection fraction (EF) using an M-mode echocardiographic image using the following formula:

EF (%) = ((LV Vol;d – LV Vol;s) / LV Vol;d) × 100, where LV volume is calculated from the M mode with (7.0 / (2.4 + LVID)) * LVID^3^ according to the operator manual of VisualSonics Vevo 2100 Imaging System.

### Histology and image analysis

For Masson’s trichrome staining, hearts were collected fixed in 4% paraformaldehyde overnight, dehydrated, paraffin-embedded, and prepared in 5-μm sections. Staining was performed according to standard procedures by the Neuropathology Core at Baylor College of Medicine. Wheat germ agglutinin (WGA) staining was performed on paraffin-embedded cross-sections of left ventricles, using tetramethylrhodamine isothiocyanate-conjugated wheat germ agglutinin (20 g/ml in PBS) (Sigma L5226), which was used to measure the cross-sectional area of cardiomyocytes. Cardiomyocyte diameter and fibrosis area were visualized with a Leica DMi8 automated fluorescence microscope and quantified, in a blinded manner, using ImageJ software version 2.0, with five microscopic fields per heart.

### Recombinant DNA and virus

Construction of Ad.U6.sgHDAC3.EF1.Cas9 was previously described ^43^. The sequence of sgHDAC3 is GTAGAAATACGCCACGGTCT. pFUGW lentiviral plasmids expressing Flag-tagged HDAC3-WT, Y298H, Δ33-70, WT-LAP2β were generously provided by Dr. Rajan Jain (University of Pennsylvania). cDNA of wild type and mutant HDAC3 were PCR amplified from corresponding Lentivirus vectors and cloned into sites between T7 promoter and bGH poly (A) signal of pENTR-CMV (Addgene, 32688) plasmids. pENTR-CMV-LAP2β-HA was generated by overlap extension PCR according to standard protocols. pAd plasmids and recombinant adenovirus vectors were then produced using the ViraPower adenoviral expression system (ThermoFisher).

### Cell culture, transfection, infection, and isotope tracing

AC16 cells were cultured in Dulbecco’s modified Eagle’s medium/Nutrient Mixture F-12 (DMEM/F12) (Fisher, 10-090-CV) containing 10% fetal bovine serum (FBS), 100 U/mL penicillin, 100 ug/mL streptomycin, and plated on 100 mm cell culture dish at 37℃ under 5% CO_2_ in a humidified incubator. Human induced pluripotent stem cells (iPSCs) were maintained in mTeSR1 medium (STEMCELL Technologies, #85850) on Matrigel-coated cell culture plates (Corning, #354248; Corning, #3506). Cells were utilized between passages 20 and 80 and passaged every 3–5 days at a confluency of 85–100% using Accutase (STEMCELL Technologies, #07920). The cardiac differentiation protocol was performed as previously described ^49^. For adenovirus infection, AC16 cells were seeded into 6-well plates prior to infection. Adenoviruses were infected into cells when they reached 60-70% confluence. Cell culture media was changed after 24 hours. AC16 cells were infected for 48-72 hours before all experiments. Control viruses are Ad-U6-GFP adenovirus with the same vector of Ad-U6-sgHDAC3. For drug treatment, cells were preincubated with Suberoylanilide hydroxamic acid (SAHA) (caymanchem, 10009929) or Entinostat (MS275) (AdooQ Bioscience, A10611) for 48 hours at different dosages before associated experiments. HEK293 cells were cultured in DMEM (high glucose, VWR 16750-074) containing 10% FBS, 100 U/mL penicillin, 100 ug/mL streptomycin, and plated on 100mm cell culture dish at 37℃ under 5% CO_2_ in humidified incubator. For transfections, HEK293 cells were seeded into 6-well plates prior to transfection. Plasmids were co-transfected into cells when they reached 60-70% confluence with jetPRIME *in vitro* DNA transfection reagent (VWR, 89129-924) according to the manufacturer’s instruction. LAP2β and HDAC3 plasmids were mixed at 1:1 molar ratio for transfection. The medium was changed after 4 hours. After transfection for 48 hours, samples were collected to perform co-immunoprecipitation. For fatty acids oxidation, AC16 cells were seeded into 6-well plates to 80%-90% confluence and were incubated in PBS with BSA-conjugated [9,10-^3^H(N)]-palmitate and carnitine for 2 h at 37 °C in the incubator. The resultant ^3^H_2_O in the incubation solution was separated from precursors using ion-exchange columns (DOWEX 1X4-400) and was measured by a scintillation counter^43^. For non-radioactive FAO assay using a fluorescence-based fatty acid oxidation assay kit from Abcam (ab222944), AC16 cells were seeded in a Matrigel-coated Costar 96 well assay plates (black wall with clear flat bottom; Corning Incorporated, USA) at a density of 5 × 10^4^ cells/well in 200 μL culture medium and allowed to adhere to the plate for 24 hours. Infection were similar with above. The assay was performed according to the manufacturer’s protocol for the baseline activity with oleate conjugated with BSA as the substrate. The oxygen consumption rate was measured with excitation 380 nm and emission 650 nm and was expressed as the initial rate of increase in the fluorescent signal, as recommended by the manufacturer.

### Immunoprecipitation, HDAC assay, and western blot

For immunoprecipitation, cells were lysed in the RIPA buffer containing 0.1% SDS, 1% NP-40, phosphatase inhibitors, and protease inhibitors. Lysates were precleared twice for one hour each with Protein G dynabeads (Fisher, 10003D) and immunoprecipitated overnight at 4 degrees with Protein G dynabeads with anti-Flag M2 Affinity Gel (Sigma, A2220-5ML) or IgG (Santa Cruz, sc-2025). Beads were collected, washed 3 times in lysis buffer, and eluted into sample buffer (Bio-Rad, 5000006) containing 0.1M DTT and 10% 2-Mercaptoethanol (Sigma, M6250). Samples were boiled for 10 minutes and analyzed by western blot. For the HDAC assay, heart tissues were lysed in RIPA lysis buffer containing 0.1% SDS, 1% NP40, 0.5% sodium deoxycholate, and phosphatase/protease inhibitors. An equal amount of total protein from each sample was subjected to immunoprecipitation with HDAC3 antibodies (Abcam 7030) followed by protein A agarose beads (Invitrogen Cat#15918014). After washing with lysis buffer, the beads were dried using an insulin syringe and mixed with the working solution containing a fluorescence-tagged acetylated peptide from the HDAC assay kit (Active Motif Cat#56200). The reaction was allowed to last for 40 min before quenching with the developing solution containing HDAC inhibitors, followed by fluorescence measurement in a plate reader. For western blot, protein lysates from different resources were resolved by Tris-glycine SDS-PAGE, transferred to PVDF membrane, and blotted with antibodies against HDAC3 (Abcam, 7030), Histone H3 (Abcam, 1791), Histone H3 acetyl K27 (H3K27ac) (Abcam, 4729), NCOR1^46^, TBLR1 (IMGENEX, IMG591), HA (Abcam, 236632), GAPDH (Cell Signaling Technology, 2118). Images were acquired using LumiQuant AC600 (Acuronbio Technology Inc).

### RT-qPCR, RNA-seq, and data processing

For RT-qPCR, total RNA was extracted using TRIzol (Sigma) and RNeasy Mini Kit (Qiagen). Reverse transcription and quantitative PCR were performed with the High Capacity RT kit, SYBR Green PCR Master Mix, and the Quant Studio 6 instrument (Life Science) using the relative quantification method with standard curves. 18S RNA was used as the housekeeping denominator. mRNA expression levels are shown relative to WT mice or Sham-treated cells. RNA-seq was performed using total RNA (n = 3 in each group). Sequencing libraries were run on the BGI MGISEQ-2000 platform to an average depth of 60 million reads per sample. The sequencing data was filtered with SOAPnuke (v1.5.2) by removing reads containing the sequencing adapter. The resultant clean reads were obtained and stored in FASTQ format and mapped to the reference genome GRCm38.p6 using HISAT2 (v2.0.4). Bowtie2 (v2.2.5) was applied to align the clean reads to the reference coding gene set, and the expression level of genes was calculated with RSEM (v1.2.12). Differential expression analysis was performed using DESeq2 (v1.4.5). A gene was considered differentially expressed if the adjusted *p*-value was < 0.05. We carried out functional annotation analysis using DAVID Bioinformatics Resources 6.7. Differentially expressed genes were used as input gene lists, and all genes expressed in the heart were used as the background. We looked for enrichment for genetic association with biological processes in Gene Ontology (GO) and KEGG pathways.

### Statistics

Results are presented as mean ± S.E.M. Differences are analyzed by two-tailed unpaired Student’s t-test for experiments with two groups and one-way analysis of variance (ANOVA) with Holm-Sidak post hoc analysis for multiple comparisons in experiments including ≥ 3 groups. All experiments were performed at least twice using distinct cohorts of mice or independent biological samples, except the RNA-seq, which was performed once. Statistical analyses were conducted using GraphPad Prism software 8.0. No statistical analysis was used to predetermine sample sizes. Instead, sample sizes were estimated based on previous publications and our previous experience required to obtain statistically significant results. The sample size for each group was indicated in the figures, figure legends, or the methods section above. Animals were excluded if they showed distress, infection, bleeding, or anorexia due to surgery or treatment. Experimental mice were randomly assigned to each experimental/control group. Investigators were blinded to the genotypes of the individual animals during experiments and results assessments. DEGs were identified by adjusting the *p*-values for multiple testing at an FDR (Benjamini Hochberg method) threshold of < 0.05.

## ACKNOWLEDGEMENT

We thank Dr. Mitchell Lazar (University of Pennsylvania) for the NS-DADm and HDAC3 loxP mouse lines, Dr. Rajan Jain (University of Pennsylvania) for LAP2β0-related lentiviral plasmids, Dr. Chris Ward at Baylor College of Medicine (BCM) Mouse Metabolism and Phenotyping Core (R01DK114356 and UM1HG006348) for echocardiography instrument, and BCM Neuropathology Core (P50103555) for histology analyses. The laboratories of the authors were supported by grants from the American Heart Association (AHA 30970064) and the National Institute of Health (NIH HL153320). The investigators were supported in part by the National Natural Science Foundation of China (82200411) and Beijing Nova Program (20240484662).

## AUTHOR CONTRIBUTION

SQ collected most of the data. CZ performed transverse aortic constriction surgery and provided training on echocardiography. WL constructed plasmids and maintained mouse lines. SS and GL helped with some of the echocardiography analysis. ZC assisted in analyzing and uploading transcriptomics data. WZ performed WGA staining. HY maintained the mouse lines. HL, HS, and ZS obtained funding. ZS conceived the study. SQ, CZ, WL, SS, GL, HS, and ZS analyzed and interpreted the data. SQ and ZS wrote the manuscript with input from the other authors.

---

**Supplemental Figure S1.**
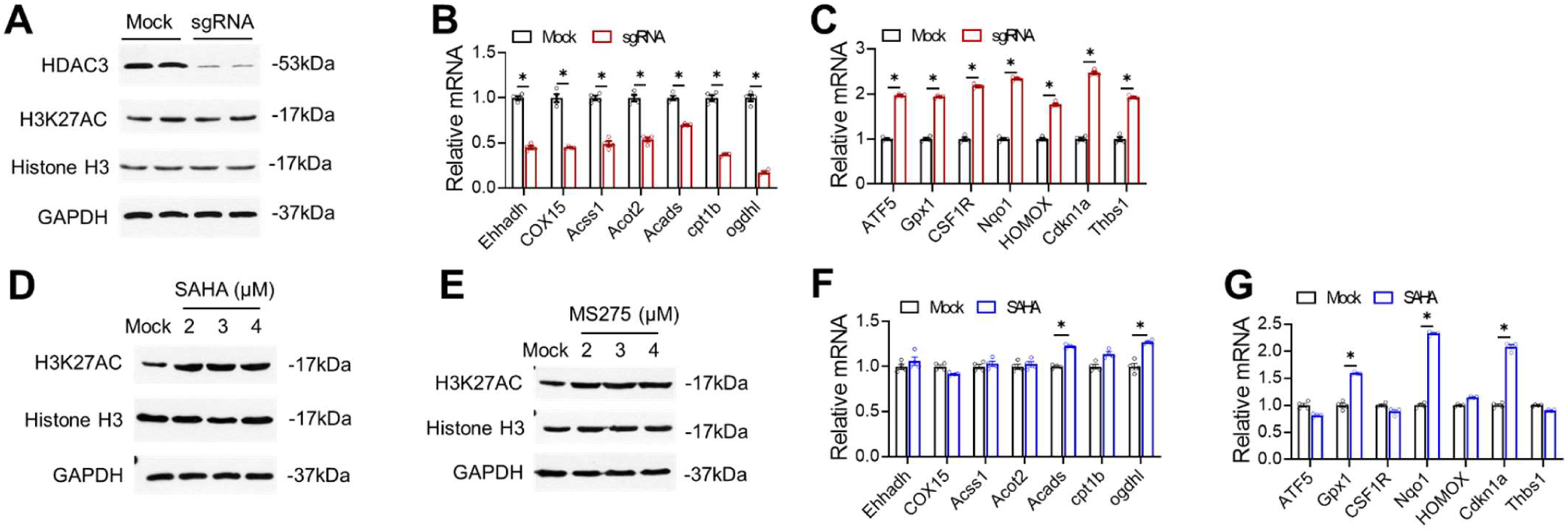
Discrete cell-autonomous effects between HDAC3 depletion and HDAC enzyme inhibition in iPSC-derived cardiomyocytes (iPSC-CM). (**A**) Western blot analysis of lysates from iPSC-CM with HDAC3 knock-down. (**B-C**) RT-qPCR analysis of the iPSC-CM with HDAC3 knock-down. n = 4. (**D-E**) Western blot analysis of lysates from iPSC-CM administrated with 2 uM MS275 or SAHA at the indicated dosages, along with vehicle control (mock). (**F-G**) RT-qPCR analysis of the iPSC-CM treated with 2 uM SAHA. All data are mean ± S.E.M. * *p* < 0.05 by t-test.

**Supplemental Figure S2.**
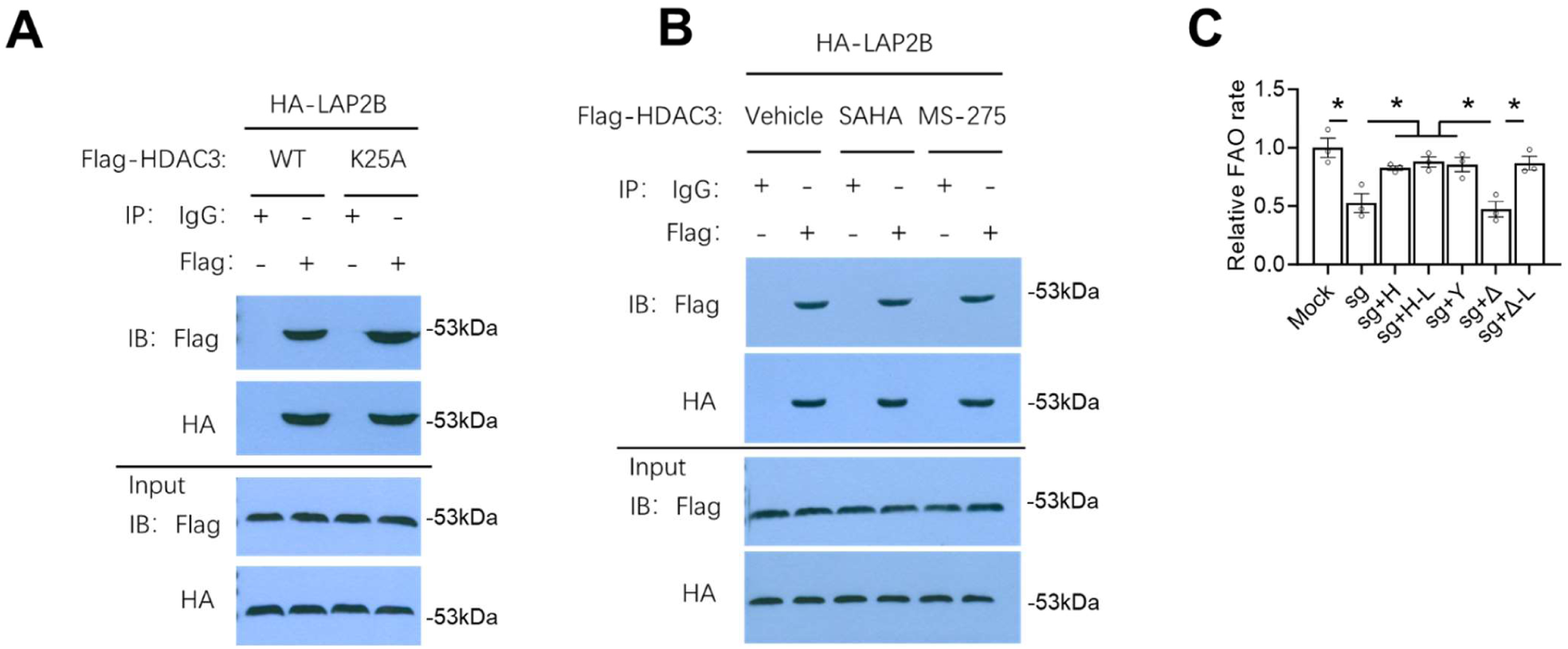
HDAC3-LAP2β interaction is not affected by HDIs or HDAC3-NCOR interaction. (**A**) Co-immunoprecipitation analysis (co-IP) with the indicated antibodies of cell lysates from HEK293 cells transfected with indicated plasmids, followed by immunoblot (IB) with the indicated antibodies. (**B**) Co-IP analysis of HEK293 cells treated with 2 uM MS275 or SAHA. (**C**) Fatty acid oxidation (FAO) assay in AC16 cells infected with the indicated adenovirus vectors. Data are mean ± S.E.M. * *p* < 0.05 by one-way ANOVA with Holm-Sidak’s post hoc.

